# Increased cortical volume without increased neuron number in heterozygous *Chd8* mutant mouse cortex

**DOI:** 10.1101/2021.01.11.426290

**Authors:** Cesar P. Canales, Samuel Frank, Jeffrey Bennett, Paris Beauregard, Pierre Lavenex, David G. Amaral, Alex. S. Nord

**Affiliations:** Center for Neuroscience, UC Davis, Davis, CA, USA.; Department of Psychiatry and Behavioral Sciences, The M.I.N.D. Institute, UC Davis, Davis, CA, USA.; Laboratory of Brain and Cognitive Development, Institute of Psychology, University of Lausanne, Switzerland.

**Keywords:** ASD, Macrocephaly, Stereology, Chd8, mouse model

## Abstract

De novo mutations in the chromatin-remodeling factor *CHD8* (Chromodomain-Helicase DNA-binding protein 8) have emerged as a key genetic risk factor for Autism Spectrum Disorder (ASD) and, more generally, neurodevelopmental disorders. Individuals with heterozygous mutations in *CHD8* typically present hallmarks of ASD with comorbid cognitive disability and macrocephaly. Knockdown or haploinsufficiency of *Chd8* in animal models has recapitulated phenotypes observed in patients, including increased head circumference and brain size. Here, we aimed to determine whether increased neuron numbers or soma size drives increased cortical volume. We performed design-based stereological analyses of cortical structure in adult male and female heterozygous *Chd8* mice and wild-type littermate controls. *Chd8* haploinsufficient male mice displayed a ~8-12% increase in cortical volume, no differences in cortical neuron number and comparable neuronal soma size. Our study reproduced previous reports of increased brain size associated with *CHD8* mutation in humans and mice and are consistent with reported sex-specific impacts of *Chd8* mutations in mice and increased burden of *CHD8* mutations in human males with ASD. These findings suggest that the nature of the cortical enlargement due to *Chd8* haploinsufficiency is complex and appears to be due to a factor other than an increased neuron number or soma size.

**Lay Summary:** We measured the size and neuron number in the neocortex in mice with heterozygous *Chd8* mutation, a model relevant to Autism Spectrum Disorder. We found an increased cortical volume in male mutants, which was not accompanied by increased neuron number or soma size. Our results indicate that the enlarged brain in *Chd8* mutant mice is complex, more evident here in males, and is due to factors other than increased neuron number.

## Introduction

Autism spectrum disorder (ASD) is a genetically heterogeneous disorder that involves the interaction of both common and rare variants and produces a broad phenotypic spectrum (Geschwind, 2009). Despite the clinical and genetic heterogeneity that typifies ASD, some anatomical features appear to be either present in many cases or so dramatically altered in some that their presence is now reasonably well replicated as characteristic of ASD among a subset of cases across a number of studies. One such finding is the tendency towards increased size of the brain during the early postnatal period causing macrocephaly (Albores-Gallo et al., 2017; Amaral et al., 2017; Bernier et al., 2014, 2016; Donovan & Basson, 2017; Li et al., 2017; Libero et al., 2016; Nordahl et al., 2011; Woodbury-Smith et al., 2017; Wu et al., 2020).

Mutations in chromatin remodeler factors (CRFs) are strongly associated with causal roles across Neurodevelopmental Disorders (NDDs) (De Rubeis et al., 2014; Gilissen et al., 2014; Iossifov et al., 2014; Nishiyama et al., 2004; O’Roak et al., 2012; O’Roak et al., 2012; Sanders et al., 2015; Vissers et al., 2016; Vogel-Ciernia & Wood, 2014). Of the top ASD risk genes ranked by enrichment of de novo case mutations, approximately 40% are CRFs (Sanders et al., 2015; Satterstrom et al., 2020). Heterozygous mutations to different CRFs cause similar phenotypes (Bögershausen & Wollnik, 2018), suggesting shared mechanisms of brain pathology. The gene *CHD8,* which encodes the CRF chromodomain helicase DNA-binding protein 8, has been implicated in genetic analyses profiling rare and *de novo* coding variation in ASD and intellectual disability (ID) (An et al., 2020; Barnard et al., 2015; De Rubeis et al., 2014; O’Roak et al., 2012; Tatton-Brown et al., 2017). Along with ASD, humans with *CHD8* mutations frequently exhibit ID, macrocephaly, craniofacial dysmorphology, and gastrointestinal dysfunction (Barnard et al., 2015; Beighley et al., 2020; Bernier et al., 2014).

The proportion of individuals with *CHD8* mutations who present macrocephaly is much higher than that seen in the general ASD population, and NDD-relevant behaviors and macrocephaly have been reported across multiple *Chd8* mutant mouse models (Barnard et al., 2015; Bernier et al., 2014; Gompers et al., 2017; Katayama et al., 2016; Platt et al., 2017a; Suetterlin et al., 2018; Zhao et al., 2018). We previously reported the impact of germline heterozygous frameshift *Chd8* mutation (*Chd8^+/del5^*) on neurodevelopment in mice (Gompers et al., 2017). *Chd8^+/del5^* mice exhibited cognitive impairment correlated with increased regional brain volume. To date, the differences underlying increased brain and cortical volume in mature mutant *Chd8* mice remain undefined. Towards advancing understanding of megalencephaly and increased brain volume associated with *Chd8* haploinsufficiency, we applied design-based stereology to characterize the neocortex of *Chd8^+/del5^* mice. Design-based stereology is now the gold-standard method to obtain reliable quantitative measurements of brain structure (Hilgetag et al., 2018; Kipp et al., 2017). We estimated cortical volume, neuron number and neuronal soma size in male and female *Chd8^+/del5^* mice and their wild-type littermates.

## Methods

### Animals

Generation of Cas9-mediated 5bp frameshift deletion in exon 5 of *Chd8* in mice was previously described (Gompers et al., 2017). Thirty-two 42-day-old mice were included in this analysis (16 brains per genotype, 16 per sex). All protocols utilized in the generation of brain histological samples were approved by the Institutional Animal Care and Use Committees (IACUC) at the University of California Davis. Mice were housed in a temperature-controlled vivarium maintained on a 12-h light–dark cycle. We made efforts to minimize pain, distress and the number of animals used in the study.

### Tissue preparation and Nissl Stain

Mice were anesthetized using Isoflurane and perfused transcardially using room temperature saline solution (0.9% NaCl) followed by 4% paraformaldehyde (PFA) in 0.1 M phosphate buffered (PB) saline using a peristaltic pump (Minipuls 3, Gilson) at 20 rpm flow rate. Brains were removed and post-fixed in 4% PFA at 4°C for 24 h. Fixed brains were cryoprotected in 50 ml of 10% and 20% glycerol solutions with 2% DMSO in 0.1 M phosphate buffer (PB; pH 7.4; for 24 and 72 hours, respectively). Brains were blocked at −2.00mm posterior to the interaural line prior to flash freezing. Sections were cut using a freezing, sliding microtome in 4 series at 60 µm (Microm HM 440E, Microm International, Germany). Every 4th section was collected in 10% formaldehyde in 0.1 M PB (pH 7.4) and post-fixed at 4°C for 1 week (Chareyron et al., 2011). Sections were rinsed twice for 1 hour each in 1ml of 0.1M PB, mounted on gelatin-coated slides, and air-dried overnight at 37°C. Defatting was performed in a mixture of chloroform/ethanol (1:1, vol) for 2 x 1 hour. The brains were rehydrated in graded alcohols; 100% ethanol, 100% ethanol, 95% ethanol, 70%, 50% and dipped in deionized water (Chareyron et al., 2011; Lavenex et al., 2009). Sections were stained 40 seconds in a 0.25% thionin solution (Fisher Scientific, Waltham, MA, cat. no. T-409), dehydrated through graded alcohols, differentiated in 95% ethanol and 5 drops of glacial acetic acid, cleared in xylene and cover-slipped with DPX (BDH Laboratories, Poole, UK) (Chareyron et al., 2011).

### Stereology

The inferior boundary of the neocortex was defined as the bottom of the agranular insular area (AIA), and the superior boundary was defined as the anterior cingulate area (ACA) or the retrosplenial area (RSA) and the white matter was then followed to the ACA/RSA border (Franklin, 2013). Neuron number, neuronal soma size, and brain volume were estimated using Stereoinvestigator V10.50 (MBF Biosciences, Williston, VT) attached to a Zeiss Imager.Z2 Vario with a Zeiss AxioCam MRc. An EC PlanNeoFluar 2.5 x objective was used to trace the outline of the neocortex and a PlanApoChromat 100x oil objective was used for cell counts and soma size determinations. The volume of the neocortex was determined using the Cavalieri method (Gundersen & Jensen, 1987; Lavenex et al., 2000; Mark J. West & Gundersen, 1990). For each mouse, 15-18 sections (480 µm apart) were analyzed with the first section chosen randomly amongst the first through the neocortex. Estimates of neuron numbers were obtained using the Optical Fractionator method (Gundersen, 1986; M. J. West et al., 1991). Since no lateralization was found in the volumetric estimates (p = 0.65), we sampled the left hemisphere of half of the mice and the right hemisphere of the other half for estimates of neuron numbers. We used the same sections that were used for volumetric estimates. We used a counting frame size of 18×18×10µm with 2 µm guard zones, and a scan grid of 480×480 µm placed at a random angle. Neurons were defined following established protocols (Gundersen, 1986; Gundersen & Jensen, 1987; Lavenex et al., 2000; Long et al., 1998; M. J. West et al., 1991; Mark J. West & Gundersen, 1990), based on: 1) dark staining, 2) a well-defined nucleolus and 3) a round, non-jagged cell body (Chareyron et al., 2011; García-Cabezas et al., 2016; Haida et al., 2019; Lauber et al., 2018; Long et al., 1998). Section thickness was measured at every counting site. The volume of neuronal somas was determined using the Nucleator method (Gundersen, 1988) during the Optical Fractionator analysis. Comparison of stereological measures (volume, neuron number, and soma size) were performed via ANOVA to examine global effects of sex and genotype and test for sex by genotype interaction, and then via t-test to compare the effect of genotype in males and females. In addition to test results, effect sizes are reported for ANOVA (partial eta squared, *ηp^2^*) and t-test (Cohen’s d). Statistical analyses were performed using SPSS (version 26), plots were generated using *ggplots2* package in R programming language (Wickham, 2009). All stereological analyses were carried out blinded to the experimental condition or sex of the animal.

## Results

Nissl-stained brain sections were used to perform design-based stereological analyses of cortical structure in *Chd8^+/del5^* and wild-type mice at postnatal day 42, an age that may represent the end of the adolescence period in mice (Spear, 2000) **(Fig 1)**. Considering cortical volume **(Fig 2a)**, there was an effect of sex (*F*(1,28) = 4.881, *p* = 0.036, *ηp^2^* = 0.148), an effect of genotype (*F*(1,28) = 6.190, *p* = 0.019, *ηp^2^* = 0.181), and an interaction between sex and genotype that neared statistical significance (*F*(1,28) = 2.956, *p* = 0.097, *ηp^2^* = 0.095). In males, cortical volume was larger in *Chd8^+/del5^* mice than in wild-type littermates (t(14) = 3.103, p = 0.008, d = 1.539). In contrast in females, cortical volume did not differ between *Chd8^+/del5^* and wild-type mice (t(14) = 0.523, p = 0.609, d = 0.249). Given the larger cortical volume in male *Chd8^+/del5^* mice, we sought to determine whether this was due to differences in neuronal number **(Fig 2b)**. There was no effect of sex (*F*(1,28) = 2.619, *p* = 0.117, *ηp^2^* = 0.086), no effect of genotype (*F*(1,28) = 0.107, *p* = 0.747, *ηp^2^* = 0.004), and no interaction between sex and genotype (*F*(1,28) = 1.968, *p* = 0.172, *ηp^2^* = 0.066). The number of cortical neurons did not differ between *Chd8^+/del5^* and wild-type littermates in either males (t(14) = 0.921, p = 0.373, d = 0.460) or females (t(14) = 1.065, p = 0.305, d = 0.532). We also sought to determine whether a larger cortical volume in male *Chd8^+/del5^* mice was associated with differences in neuronal soma volume **(Fig 2c)**. There was an effect of sex (*F*(1,28) = 4.611, *p* = 0.041, *ηp^2^* = 0.141; males > females), but no effect of genotype (*F*(1,28) = 1.705, *p* = 0.202, *ηp^2^* = 0.057) or interaction between sex and genotype (*F*(1,28) = 1.121, *p* = 0.299, *ηp^2^* = 0.038). Neuronal soma size did not differ between *Chd8^+/del5^* and wild-type littermates in either males (t(14) = 0.242, p = 0.812, d = 0.120) or females (t(14) = 1.375, p = 0.191, d = 0.687).

**Figure 1:**
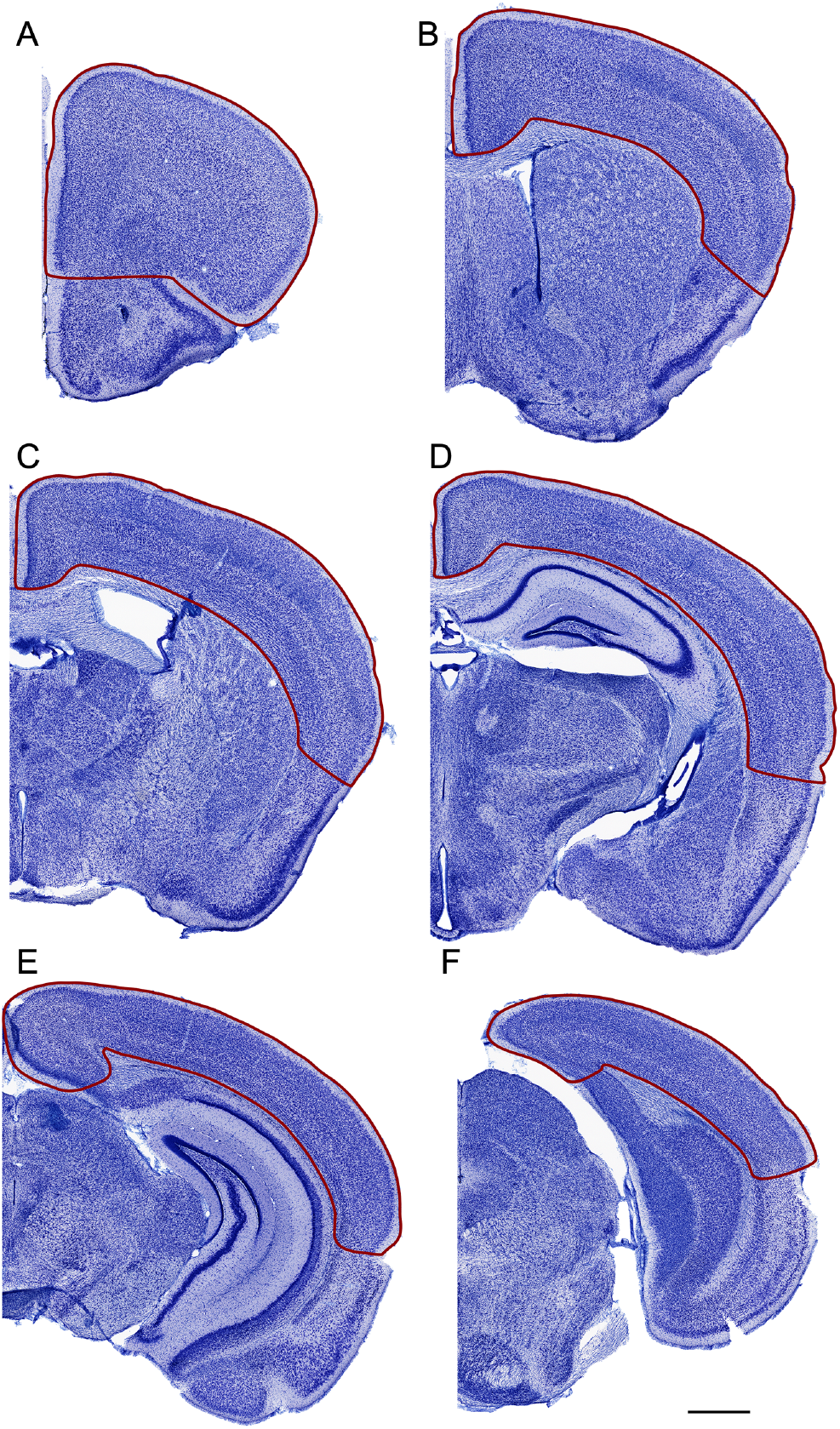
Nissl-stained coronal sections through the mouse brain (arranged from rostral A, to caudal F) illustrating the portion of the neocortex that was evaluated in this study. Not all levels are shown and the analyzed neocortex extended to the rostral and caudal poles of the brain. Before the study was started, a template of cortical areas was designed that was based on identifiable neuroanatomical landmarks. The template was used to demarcate the extent of analyzed neocortex in all experimental animals. The analysis was carried out blinded to the experimental condition or sex of the animal. Calibration bar equals 1 mm.

**Figure 2:**
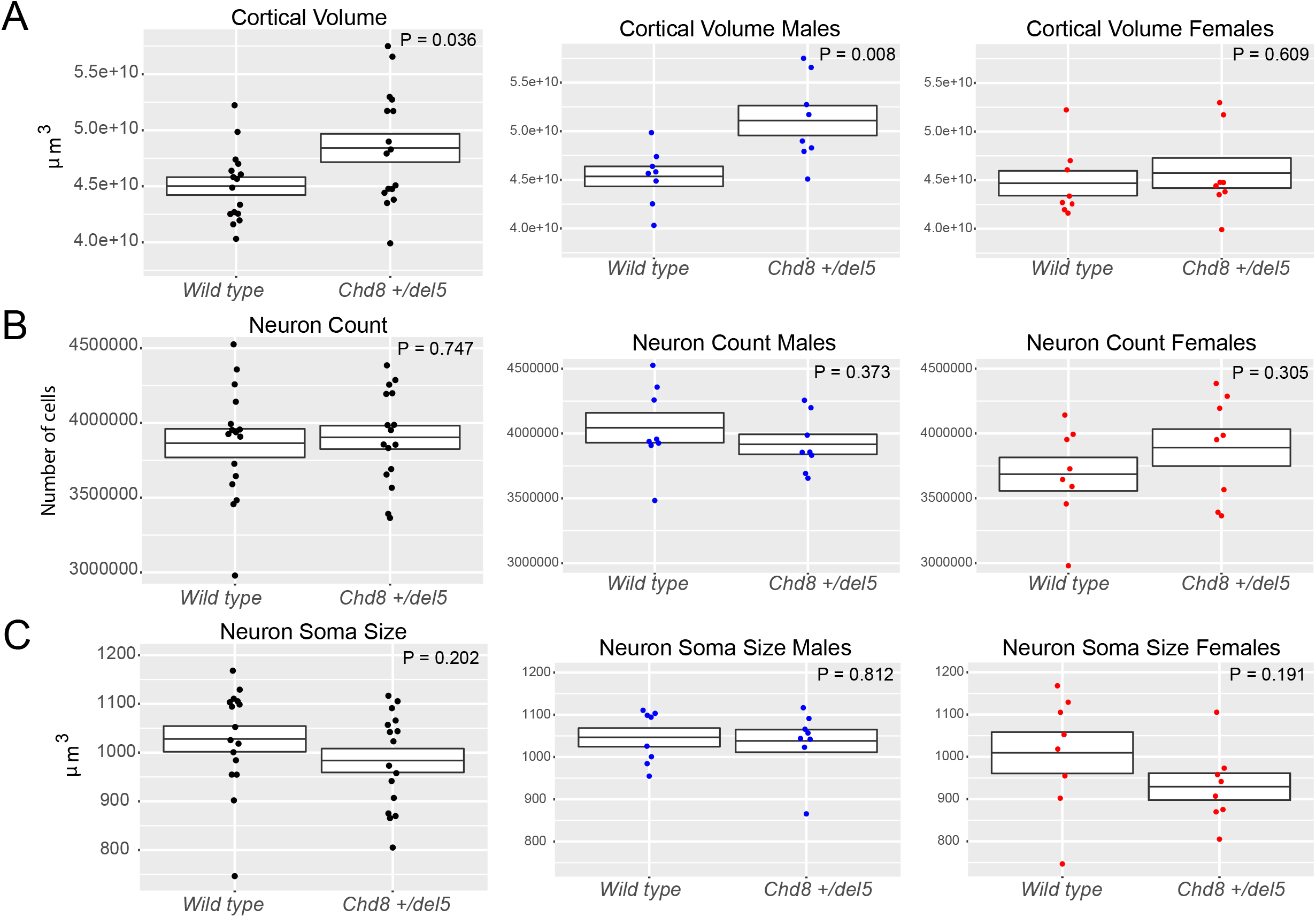
Increased cortical volume without increased neuron number in heterozygous Chd8 mutant mice. Stereological estimations of Cortical volume (a), Neuron count (b), and Neuron Soma Size (c) measured on Nissl-stained sections from *Chd8^+/del5^* mice and wild type littermates. P values resulting from t-test comparisons of the effect of genotype and sex stratified dispersion plots are shown. All data expressed as mean ± SEM.

## Discussion

Germline heterozygous *CHD8* mutations in humans, mice, and zebrafish all have been reported to produce macrocephaly/megalencephaly (Barnard et al., 2015; Beighley et al., 2020; Bernier et al., 2014; Gompers et al., 2017; Katayama et al., 2016; Platt et al., 2017b; Suetterlin et al., 2018; Sugathan et al., 2014; Zhao et al., 2018). As *Chd8* haploinsufficiency has also been tied to altered neurogenesis in vitro and in vivo (Cotney et al., 2015; Durak et al., 2016; Gompers et al., 2017; Suetterlin et al., 2018), one possible explanation for a larger brain size is an increased proliferation and greater numbers of neurons. Using design-based stereology, we demonstrated a larger cortical volume in male *Chd8^+/del5^* mice. Our findings further indicate that the underlying cause of this difference, as observed in male *Chd8^+/del5^* mice here, is not an increase in neuron number or neuronal soma size in the neocortex.

Recent comparisons of *CHD8* mutations in ASD case populations indicate a higher number of male carriers, suggesting that there are sex-specific differences in susceptibility and phenotype (An et al., 2020; Bernier et al., 2016; T. Wang et al., 2016). One study of *Chd8* mutant mice similarly reported sex differences in transcriptomic and behavioral phenotypes (Jung et al., 2018). Our findings add to the evidence that males may be differentially impacted by *CHD8* mutations, as we only observed increased cortical volume in male mice in the current study. However, the literature indicates that both male and female ASD cases with *CHD8* mutations demonstrate consistent clinical phenotypes, including macrocephaly (An et al., 2020; Barnard et al., 2015; Beighley et al., 2020; Bernier et al., 2014; Douzgou et al., 2019; Li et al., 2017; T. Wang et al., 2016; Wu et al., 2020). Additionally, publications of *Chd8* mouse models, including earlier work on this line, reported relevant phenotypes in both males and females (Gompers et al., 2017; Suetterlin et al., 2018; Zhao et al., 2018). The specific *Chd8* loss-of-function mutation characterized here occurs on the C57Cl/6N inbred line, and differences across mutant alleles and inbred mouse genetic backgrounds could influence phenotype penetrance and expressivity. Future studies that are well-powered to test for sex-specific differences and incorporate genetic background are needed to determine differences in penetrance and expressivity of NDD-relevant molecular, neuroanatomical, and behavioral phenotypes in *Chd8* mouse models.

We previously reported increased embryonic neurogenesis via Edu labeling and analysis of neural progenitor and postmitotic populations at E14.5 in this same *Chd8* mutant mouse line (Gompers et al., 2017). Other studies have similarly found evidence for macrocephaly, altered neural proliferation, and altered regulation of cell cycle genes (Durak et al., 2016; Marie et al., 2018; Suetterlin et al., 2018; Sugathan et al., 2014; P. Wang et al., 2017; Xu et al., 2018).

However, the findings here suggest that any altered proliferative output during embryonic corticogenesis does not directly explain lasting postnatal increase in cortical volume in *Chd8^+/del5^* mice. A study of *Chd8* knockdown via in utero electroporation of siRNA found decreased neurogenesis at E16 (Durak et al., 2016), and increased or decreased proliferation could be transient or compensated via increased neuronal apoptosis, resulting in no overall change in total neuron numbers (Chen et al., 2015; Naruse & Keino, 1995). Factors independent of neurogenesis could contribute to increased cortical volume in mature mice, including changes in neuronal dendritic morphology, and altered number or morphology of glial cells (Matta et al., 2019; Schumann & Nordahl, 2011; Winden et al., 2015). Heterozygous mutant *Pten* mice, another ASD risk gene associated with macrocephaly, exhibit increased neurogenesis and a greater number of neurons in early postnatal brain, but change to excess glial cells in the mature brain (Chen et al., 2015). Further, changes in brain volume are dynamic across *Pten* mutant mice and with differential impacts across brain regions (Chen et al., 2015; Clipperton-Allen et al., 2019). While mutation to *Chd8* and *Pten* represent distinct models, mutations in both lead to comorbid ASD and macrocephaly and Chd8 has been implicated in *Pten* signaling pathways (Sugathan et al., 2014; P. Wang et al., 2017). Future efforts will be needed to test for the dynamic changes in corticogenesis and relevant neuronal populations across development and for differences in populations of non-neuronal cells as a factor in *Chd8*-associated macrocephaly.

## Acknowledgements

This work and ASN and CPC were supported by NIH NIMH (MH120513-02) and NIGMS (R35 GM119831-05). CPC is recipient of NIMH Institutional National Research Service Award Autism Research Training Program fellowship (T32 MH073124-16). This research was also supported by NIH Grants 1R01MH121447-01 and 1R01MH103371 to DGA and Swiss National Science Foundation grant 310030_143956 to PL. Authors declare no conflict of interest.

